# *Salmonella* Infantis, the emerging human multidrug resistant pathogen – a One Health perspective

**DOI:** 10.1101/2023.07.28.549231

**Authors:** Jennifer Mattock, Marie Anne Chattaway, Hassan Hartman, Timothy J. Dallman, Anthony M. Smith, Karen Keddy, Liljana Petrovska, Emma J. Manners, Sanelisiwe T. Duze, Shannon Smouse, Nomsa Tau, Ruth Timme, Dave J. Baker, Alison E. Mather, John Wain, Gemma C. Langridge

## Abstract

*Salmonell*a Infantis presents an ever-increasing threat to public health due to its spread throughout many countries and association with high levels of antimicrobial resistance (AMR). Whole genome sequences of 5,284 *S*. Infantis strains from 74 countries, isolated between 1989 and 2020 from a wide variety of sources including humans, animals, and food, were analysed to compare genetic phylogeny, AMR determinants and plasmid presence.

The global *S*. Infantis population structure diverged into three clusters: a North American cluster, European cluster and a global cluster. The levels of AMR varied between the *S*. Infantis clusters and by isolation source; 73% of poultry isolates had multidrug resistance (MDR) compared to 35% of human isolates. This correlated with plasmid of emerging *S*. Infantis (pESI) presence; 71% of poultry isolates contained pESI versus 32% of human isolates. This provides important information for public health teams engaged in reducing the spread of this pathogen.

## Introduction

Non-typhoidal *Salmonella* (NTS) infections place a large burden on public health, with an estimated 79 million cases of foodborne NTS infection occurring in 2010 (1). *Salmonella enterica* subspecies *enterica* serovar Infantis (*S*. Infantis) is becoming an increasingly prevalent serovar globally. An increase in human infections between 2001 and 2016 of 167% was observed in the USA (2), and in EU member states, it is the predominant serovar isolated from broiler flocks and broiler meat, accounting for 56.7% of *Salmonella* isolates from broiler meat in 2018 (3,4). Higher levels have been observed in Japan at 72.2% of isolates from ground chicken, and 84% of broilers in Ecuador (5,6).

AMR in *S*. Infantis varies by location; in South Africa only 13.4% of 387 *S*. Infantis isolates from humans had AMR (7). Conversely, in EU member states in 2016, 70% of *S*. Infantis isolates from broiler meat harboured MDR (8). Of particular concern is the emergence of extended beta-lactamases (ESBLs), such as the bla_CTX-M-65_ gene which has been reported in *S*. Infantis from countries such as Ecuador, Peru, Switzerland, UK and USA (9– 13). pESI has been found to be responsible for these high levels of AMR as it confers resistance to trimethoprim, streptomycin, sulfamethoxazole and tetracycline; ESBLs have also been found to be carried by some pESI variants (10,11,14). Originally identified in Israel, pESI-like plasmids have since been reported in multiple countries (14–19).

*S*. Infantis has a polyphyletic population structure comprising two eBurst Groups (eBG), eBG31 and eBG297 which differ by five to seven Multi-Locus Sequence Typing (MLST) alleles (16). The dominant eBG (single locus variants around a central ST) globally is eBG31; eBG297 comprised just 0.7% of *S*. Infantis isolates on Enterobase on 09.08.21 (20). However, higher levels (32%) of eBG297 have been reported in South Africa (7).

The population structure of *S*. Infantis has been studied on a limited scale; whole genome sequencing (WGS) analysis of 100 *S*. Infantis isolated from multiple continents and sources found no clustering by geographic location (16). *S*. Infantis isolates were found to cluster by isolation source from Japanese human and chicken meat samples (21), by bla_CTX-M-65_ presence in USA and Italian human and animal strains (11), and by pESI presence in human and poultry isolates from Switzerland (10).

Whilst MDR *S*. Infantis is an emerging public health concern, no large-scale population structure study of this pathogen has been performed. As eBG297 isolates have been analysed in depth (7) our aim was to determine the global population structure of eBG31 from a One Health perspective, investigating whether the population structure is associated with isolation source, location, MDR or pESI presence.

### Methods

Our eBG31 collection contained 5284 isolates, sourced from UKHSA, NICD, APHA, GenBank and Enterobase (20), detailed methods are available in Supplementary information. The collection included strains isolated from 74 countries and spanned four decades, including strains isolated between 1989 and 2020. The isolates were grouped into the following sources: animal feed, human, environmental, food, other animals, poultry, poultry products and unknown.

The whole genome consensus FASTAs were grouped into clusters where all sequences in each cluster were less than ‘n’ SNPs from another member. A core SNP phylogeny of representatives of 25SNP clusters was generated; clusters were identified using fastbaps and treedater used to date the phylogeny (22–25). AMR determinants and plasmid presence were screened for in all eBG31 sequences as described in Mattock *et al*., 2022 (7).

## Results

### Demographics

Isolates from a multitude of sources were included in the eBG31 collection. The majority, 60% (3150/5284) were isolated from humans and associated with either non-invasive infections, such as samples from stool and urine, or invasive infection with isolates from blood and cerebrospinal fluid. Six percent (300/5284) of the isolates were from poultry and 13% (684/5284) from poultry products which included samples from poultry meat, eggs and processed meals containing poultry meat. Isolates from other animals comprised 6% (321/5284) of the collection; 7% (390/5284) were from food; 1% (74/5284) from animal feed and 5% (268/5284) were environmental isolates including water, farm swabs and soil.

## Ninety-seven isolates had no stated isolation source

The number of isolates increased temporally until 2017; this was due to isolates from public databases being included until February 2018 (Figure S1), only strains isolated by the UKHSA were included following this. As shown in Figure 1, the United States contributed the largest number of isolates (n=2719). When split by continent 54% (2861/5284) of the strains were from North America; 31% (1642/5284) from Europe; 6% (316/5284) from Africa; 6% (312/5284) from Asia; 2% (128/5284) from South America and 0.47% (25/5284) were unknown.

**Figure 1.**
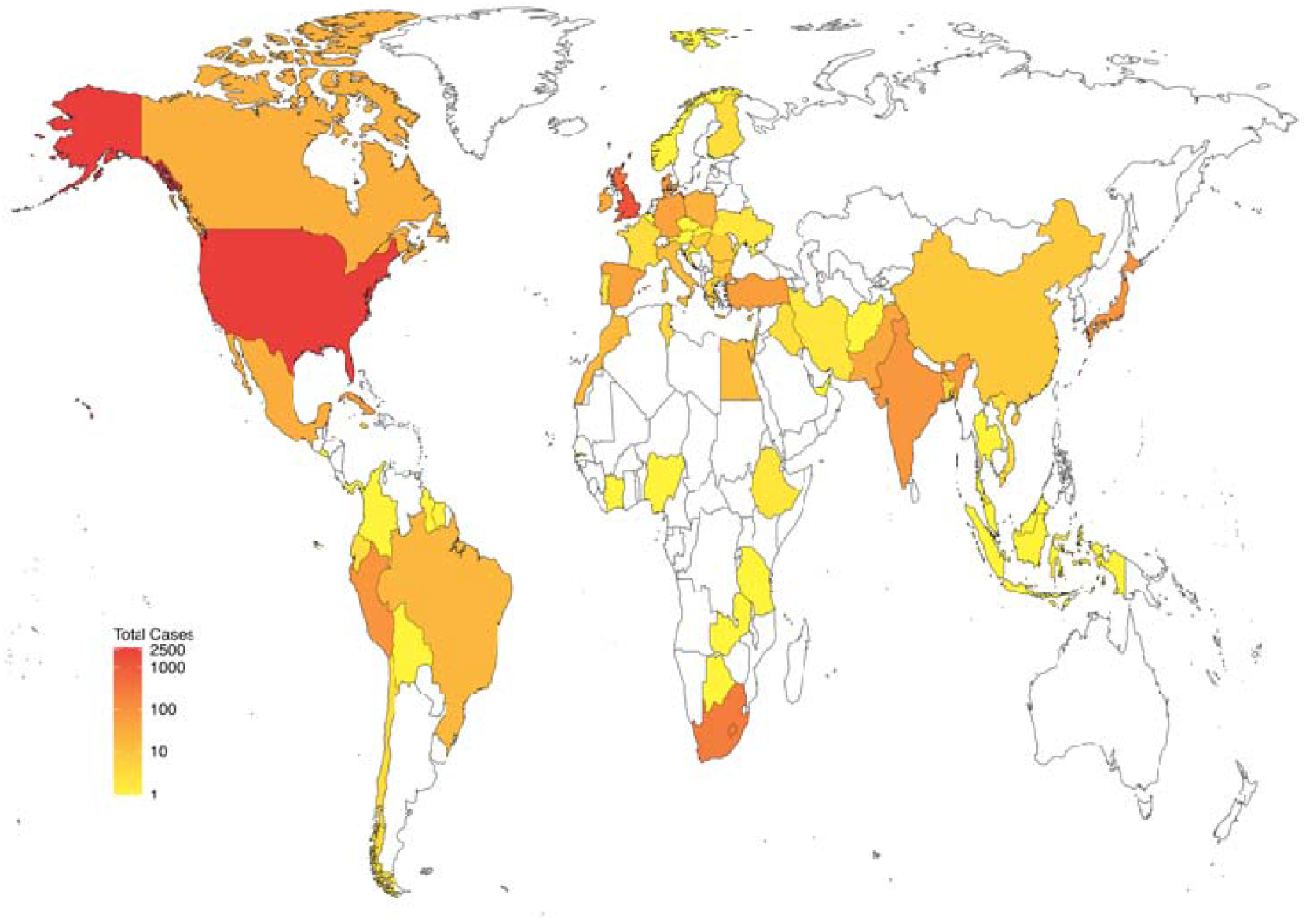
Frequency of *S*. Infantis from each country Heatmap of the number of *S*. Infantis strains included in the dataset from each country

## Population Structure

At the ST level the vast majority of eBG31 isolates, 99% (5205/5284), belonged to ST32. The second most common was ST2283, with 36 isolates, all from Europe, 17 from humans, 10 from other animals and five from food. This was followed by ST2146 with 26 isolates, all from North America, 22 of which were from environmental samples, three from food and one from a clinical sample. The 13 remaining STs were found in three or fewer isolates.

Three 250SNP clusters were present, one containing just SRR8114924. There were 408 50SNP clusters, 1288 25SNP clusters, 2876 10SNP clusters and 3917 5SNP clusters. Figure 2 is a core SNP maximum likelihood phylogeny of a member of each 25SNP cluster, representing 5283 eBG31 isolates; it is available as a cladogram in Figure S2. Bayesian hierarchical clustering identified three clusters: Cluster A contained 348 sequences, representing 1624 isolates (Figure 2, blue); Cluster B had 831 sequences, representing 3283 (Figure 2, pink) and Cluster C, which diverged from within Cluster B, contained 109 sequences, representing 376 (Figure 2, purple). When annotated by ST the phylogeny was dominated by ST32 (Figure S3); 99% (1269/1288) of the 25SNP clusters were exclusively ST32. The three 25SNP clusters containing the ST2283 isolates clustered together in Cluster C and another three clusters comprising the ST3815 isolates clustered in Cluster A.

**Figure 2.**
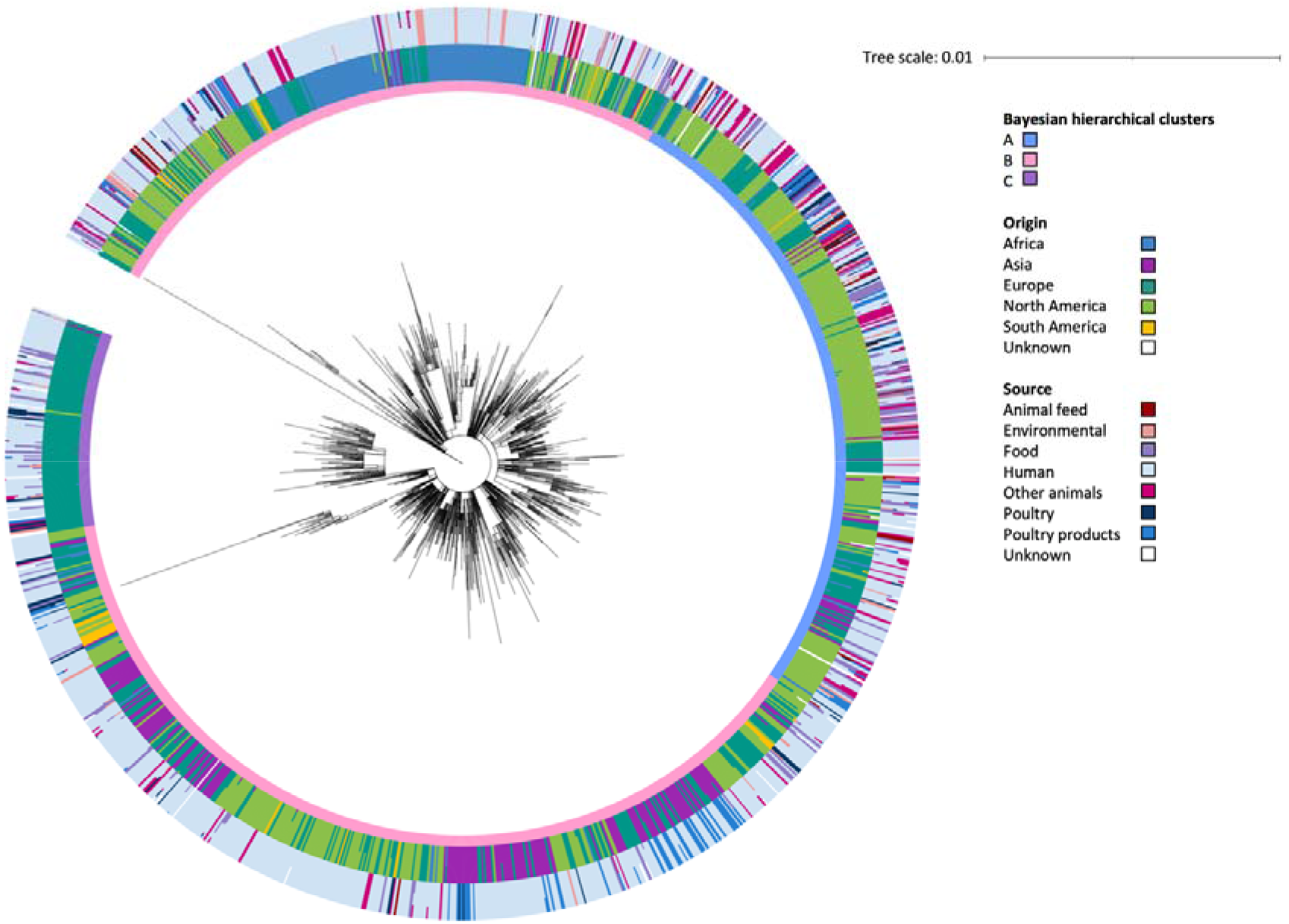
Maximum likelihood phylogeny of *S*. Infantis Core SNP maximum likelihood phylogeny of 1288 representatives of 5283 *S*. Infantis isolates. The inner ring around the phylogeny is annotated with the Bayesian hierarchical clusters found by fastbaps. The outer rings show the percentage of isolates in each 25SNP cluster that were from each continent and source. Bayesian hierarchical clusters: A, 348 representatives of 1624 isolates; B, 831 representatives of 3283 isolates and C 109 representatives of 376 isolates Origin: Africa (n=316), Asia (n=312), Europe (n=1641), North America (n=2861), South America (n=128) and unknown (n=25) Source: animal feed (n=74), environmental (n=268), food (n=390), human (n=3149), other animals (n=321), poultry (n=300), poultry products (n=684) and unknown (n=97)

In contrast to previous reports a geographical signal was visible in the clustering of isolates in the phylogeny. Cluster A was mainly comprised of North American isolates, with 74% (1203/1624) from that continent, 19% from Europe (312/1624) and less than 5% of isolates from Asia, Africa or South America. Cluster B also contained a large proportion of North American isolates (50%, 1657/3238) but higher proportions of isolates from all other continents were observed at 9% (294/3238), 7% (240/3238), 29% (959/3238) and 4% (124/3238) from Africa, Asia, Europe and South America respectively. Conversely, Cluster C was almost exclusively comprised of European isolates (98%, 370/376); the remaining isolates were from North America, Asia and Africa.

The predominant isolation source in each of the clusters was humans, comprising 52% (844/1624), 62% (2035/3283), and 72% (270/376) of Clusters A, B and C respectively. Isolates from environmental sources encompassed 7% (112/1624) of Cluster A, 4% (144/3238) of Cluster B and 3% (12/376) of Cluster C. Poultry isolates were most often found in Cluster B where they made up 7% (241/3283) of isolates; 3% (51/1624) of Cluster A and 2% (8/376) of Cluster C were from poultry sources. Isolates from poultry products were also most frequently observed in Cluster B at 16% (528/3283); 9% (153/1624) of Cluster A and 1% (3/376) of Cluster C were from poultry products. Thirteen percent of the Cluster C isolates (51/376) were from other food, 11% (187/1624) of Cluster A and 5% (152/3283) of Cluster B.

Minimal clustering by year was observed in the phylogeny (Figure S4). The most common year range in each cluster was 2016-2020, representing 48% (785/1624), 51% (1664/3238) and 67% (253/376) of Cluster A, B and C respectively. The earliest date of isolation varied, Cluster B was the oldest, with an isolate from 1989. Cluster A’s oldest isolates were from 1996; Cluster C appeared more recently with its oldest isolate being from 2007. The time of the most recent common ancestor was estimated to be 1946, with Cluster A diverging in 1982 and Cluster C in 1987.

The pairwise nucleotide distance was calculated between each whole genome consensus FASTA to show the diversity within and between groups of isolates. When comparing within isolation sources, strains from poultry products had the lowest median pairwise nucleotide distance at 145 (range 0-1249) and human isolates the largest at 241 (range 0-2060). The median pairwise nucleotide distance between human isolates and other sources ranged from 215 with environmental isolates (range 1-2004) to 259 with poultry isolates (range 0-2039). Poultry isolates had a similar median nucleotide distance to environmental isolates (251, range 4-1822); a larger distance was observed between poultry isolates and those from other animals (310, range 4-1817). The largest median pairwise nucleotide distance between source groups was poultry and animal feed at 334 (range 5-1614). Low median pairwise nucleotide distances were observed within isolates from South America (44, range 0-920) and North America (159, range 0-1883); the largest distance within isolates from a continent was Africa at 492 (range 0-1885). African isolates also had the largest median pairwise nucleotide distances to all other continents, from 395 (range 15-2059) with North America to 422 (range 23-1639) with South America.

### AMR in *S*. Infantis

In this collection 44% (2327/5284) of the isolates contained at least one AMR gene, most of these, 40% (2101/5284) had MDR. The association between population structure and AMR is illustrated in Figure 3. AMR genes were identified in isolates throughout the phylogeny; 46.7% (602/1288) of the 25SNP clusters contained an isolate with AMR genes. Some 25SNP clusters contained large numbers of isolates with AMR; such as one in Cluster B which contained 734 isolates of which 727 had MDR. A common resistance profile was visible in 25SNP clusters across the phylogeny: 27.7% (357/1288) contained an isolate with resistance to aminoglycosides, fluoroquinolones, sulphonamides and tetracyclines (AFST). Of these, 56.9% (203/357) had an isolate with putative trimethoprim resistance and 99.7% (356/357) contained a mutation in the quinolone resistance determining region (QRDR). The proportion of isolates with this resistance profile varied between the clusters; from 1% (16/1624) in Cluster A to 43.5% (1407/3283) in Cluster B and 84% (316/376) in Cluster C. The amount of MDR differed between the clusters: 7% (114/1624) of isolates from Cluster A had MDR; 50.5% (1657/3283) and 87.8% (330/376) from Clusters B and C respectively. A higher proportion of isolates were positive for ESBLs in Cluster B (20.5%, 672/3283) than in Clusters A (0.2%, 4/1624) and C (0%).

**Figure 3.**
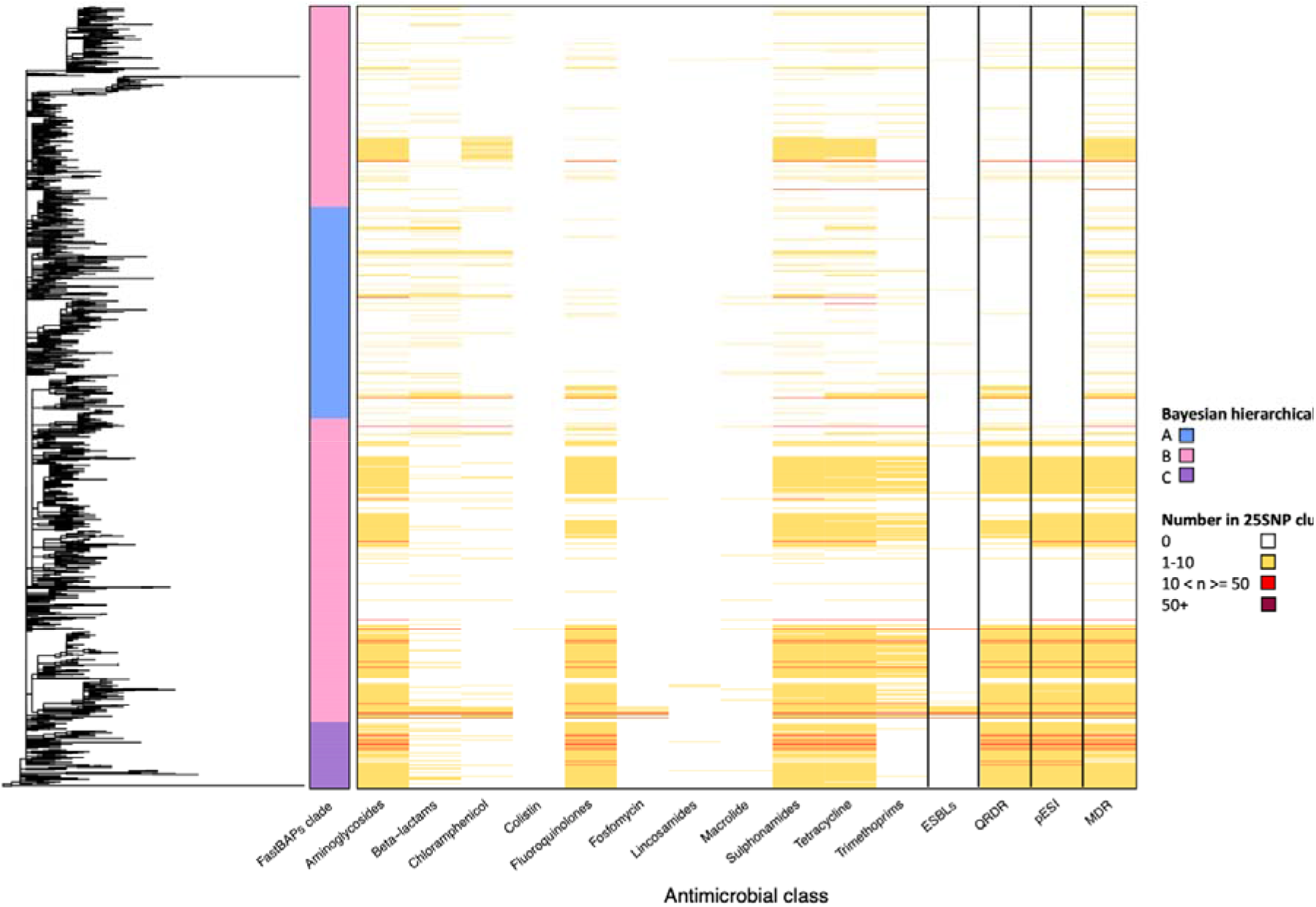
Heatmap of AMR and pESI in the *S*. Infantis phylogeny Heatmap showing the number of isolates in each 25SNP representative cluster (n=1288) of the eBG31 maximum likelihood phylogeny with genes conferring resistance to aminoglycosides, beta-lactams, chloramphenicol, colistin, fluoroquinolones, fosfomycin, lincosamides, macrolides, sulphonamides, tetracyclines, trimethoprim and ESBLs. Fastbaps cluster and the number of isolates with MDR, mutations in the QRDR conferring resistance to fluoroquinolones and pESI presence are also shown.

Variation in the distribution of AMR was observed between isolation sources. Isolates from animal feed contained the lowest amount of AMR, with 4% (3/74) predicted to have the AFST resistance profile. Higher levels of AMR were predicted in the human isolates, 29% (913/3150) had the AFST resistance profile and 22% (680/3150) had trimethoprim resistance genes. Substantially more AMR was present in the poultry and poultry product isolates: 61% (183/300) and 64% (438/684) respectively had the AFST resistance profile; beta-lactam and chloramphenicol resistance was more common in the poultry product isolates (43% (296/684) and 44% (302/684)) than in the poultry isolates (21% (64/300) and 22% (65/300)). ESBLs were identified in 10% (312/3150) of the human isolates, 19% (58/300) of the poultry isolates and 40% (272/684) of the isolates from poultry products. MDR was observed in 4% (3/74) of isolates from animal feed, 14% (38/268) of environmental isolates, 32% (123/390) of food isolates, 35% (1115/3150) of human isolates, 21% (67/321) of isolates from other animals, 73% (218/300) of poultry isolates and 73% (501/684) of poultry product isolates.

The AMR profiles also varied by the continent of isolation. The lowest levels of AMR were observed in isolates from Africa, of which 4% (14/316) had the AFST resistance profile; compared to 27% (763/2861) of North American isolates; 42% (692/1642) of European isolates, 55% (172/312) of Asian isolates and 76% (97/128) of South American isolates. ESBLs were present in 0.3% (1/316), 3% (9/312), 4% (65/1642), 19% (531/2861) and 55% (70/128) of isolates from Africa, Asia, Europe, North America and South America respectively. The proportion of isolates with MDR were 20% (63/316), 31% (891/2861), 48% (793/1642), 80% (248/312) and 81% (104/128) from Africa, North America, Europe, Asia and South America respectively.

The proportion of isolates with AMR fluctuated throughout the study period, with an upwards trend in the last fifteen years of the collection period (Figure S5). The earliest isolate in the collection, from 1989, was predicted to be resistant to six antimicrobial classes. AMR to aminoglycosides, sulphonamides and tetracyclines were consistently the most common and appeared to follow a similar trend; following 2013 similar levels of AMR to fluoroquinolones were also present.

### Plasmids in *S*. Infantis

As observed with AMR, under 50% (47%, 2502/5284) of *S*. Infantis isolates contained a plasmid. Some of the most common types included IncA/C (n=103), IncI1 (n=251) and IncX1 (n=65). As expected, pESI was the prevailing plasmid type, present in 36% (1912/5284) of the *S*. Infantis isolates. Low levels of IncA/C were observed in all isolation sources but animal feed, at most 4% (14/321) of other animal isolates, 2% of both human (64/3150) and poultry product (15/684) isolates and 0.6% (2/300) of poultry isolates. Incl1 was most common in other animal isolates at 9% (28/321) and IncX1 in human isolates (2%, 52/3150). IncI1 positive isolates were found in all continents throughout the duration of this project and mainly from humans (n=184); IncX1 was observed in Asia, Europe and North America.

pESI presence was observed in 71% (213/300) of poultry isolates, 71% (486/684) of isolates from poultry products, 32% (992/3150) of human isolates, 4% (3/74) of animal feed isolates, 10% (31/321) of isolates from other animals, 31% (120/390) of food isolates and 11% (30/268) of the environmental isolates. Presence also varied by geographic location: the lowest proportion of pESI positive isolates was from Africa at 4% (12/317); followed by North America at 28% (808/2861); Europe at 47% (770/1642); Asia at 71% (222/311) and South America at 77.3% (99/128). The earliest isolation of pESI in this collection was in four human isolates from Japan in 1999 (DRR022718, DRR022719, DRR022720, DRR022754). Figure 3 shows that whilst the common resistance profile AFST was distributed throughout the eBG31 phylogeny, Cluster A was lacking any pESI; the majority of Cluster C contained pESI (99.7%, 375/376) and 46.8% (1537/3283) of Cluster B.

## Discussion

In our large core SNP analysis of *S*. Infantis we determined that the global population structure of eBG31 is comprised of three clusters with varying isolation sources and levels of AMR. As observed previously the dominant ST in eBG31 is ST32, comprising 99% of the isolates (26–29). The other STs weren’t observed in multiple continents, suggesting that these STs have emerged in specific areas but not spread globally. A strong geographic signal was identified in the eBG31 phylogeny (Figure 2), we hypothesise Cluster B to be an ancestor of the two other clusters -containing more diversity (genetically and geographically) with isolates from all continents, we therefore designate this the global *S*. Infantis cluster. Cluster A, estimated to have diverged from Cluster B in 1982, was mainly comprised of isolates from North America and hence is named the North American cluster. Cluster C, which diverged from Cluster B in 1987 was dominated by isolates from Europe and is thus named the European cluster. This differs, perhaps due to our increased number of isolates, from Gymoese *et al*., 2019 who found no geographical signal when examining isolates from five continents and Alba *et al*., 2020 who reported little clustering by location or source (16,30). Some clustering by country of isolation was described in Acar *et al*., 2019; however as this was between isolates from the same region in Turkey, the contribution to global clustering was not clear (31). Nucleotide distances relative to the reference showed that the African eBG31 isolates were both the most diverse and the most distant to isolates from other continents; as our previous work identified that an increased proportion of isolates belonged to eBG297, this work concurs that the African *S*. Infantis population differs to that observed elsewhere (7).

As the majority of eBG31 isolates were from human sources the phylogeny was dominated by this source. Cluster C in particular contained lower numbers of environmental, poultry and poultry product isolates; this is possibly due to bias in data sampling as the cluster contains strains from UKHSA that were isolated after the cut-off for inclusion from Enterobase; the majority of these were from humans as environmental sampling tends to only be performed in association with an outbreak. While Cluster C contained strains isolated as early as 2007 it contained a lot of the newer strains, isolated between 2018 and 2020; this could represent an emerging clade of *S*. Infantis. The nucleotide distance between source groups, relative to the reference, identified the least diversity in the poultry product isolates and the greatest diversity between the poultry/poultry products and animal feed or other animal genomes. This could indicate a reduction of adaptation in poultry hosts and suggest that the different niches have encouraged adaptation; the reduced distance between poultry isolates could however be attributed to the large number of North American poultry isolates reducing the median, for example 519/734 of strains in the 25SNP cluster with the representative SRR2537092 were from poultry and 724 isolates in that cluster were from North America.

As described in many other studies, the *S*. Infantis in this project were associated with high levels of putative antimicrobial resistance (8,18,32,33). MDR genotypes were detected in 40% of the *S*. Infantis isolates, with a marked presence in isolates from poultry and poultry products, where 73% had MDR. Comparatively just 14% of environmental isolates, 21% of isolates from other animals, 32% of food isolates and 35% of human isolates had MDR (Table S1); this could indicate that the source of human infection is more often environmental sources. Whilst this has been suggested in Slovenia, where most broiler isolates clustered separately from human isolates (29), many incidences of human outbreaks associated with poultry have been reported (21,34,35); both environmental and poultry sources could be contributing to human cases but it’s also possible that the pESI positive strain circulating in the poultry industry has a selective disadvantage to causing infection in humans so these strains are observed less frequently.

The levels of AMR varied by continent, the lowest levels were observed in Africa (20% with MDR) and the highest in South America (81%); the latter concurs with other reports which found all but one isolate tested in Ecuador having MDR and multiple drug resistance profiles observed in *S*. Infantis from Chile (35,36). AMR fluctuated temporally, increasing in the last 15 years of the collection period. The proportion of isolates with resistance to aminoglycoside, sulphonamides and tetracyclines had a very similar trend, joined by fluoroquinolones following 2013. This could be attributed to pESI which carries resistance genes for these antimicrobials. Similarly, Figure 3 illustrates an association between pESI presence and resistance to aminoglycosides, fluoroquinolones, sulphonamides, tetracyclines, MDR and mutations in the QRDR. As the pESI backbone has been confirmed to carry *aadA1, sul1* and *tetA* (14,19) this illustrates how strong a driver of AMR pESI is in the *S*. Infantis population, agreeing with Alba *et al*., 2020 who suggested that pESI acquisition could be the decisive factor in the spread of the serovar throughout Europe (30). Interestingly, whilst Cluster C, the clade we suspect is an emerging dominant strain in Europe, is dominated by AMR and pESI; Cluster A, the North American cluster, with relatively low levels of MDR, 7%, was lacking any pESI positive isolates. pESI was present in 808 North American isolates in this dataset, they just belonged to either Cluster B or C. This concurs with previous research that reported two clades within the USA *S*. Infantis population, one without and the other with pESI and suggests that two groups of *S*. Infantis are circulating in North America; one associated with MDR and pESI and the other endemic to North America and not carrying pESI (37). pESI-like plasmids have recently been identified in *S*. Agona, *S*. Muenchen, *S*. Schwarzengrund and *S*. Senftenberg; the increased virulence of pESI positive isolates (38) and transmissibility of this plasmid within the *S*. Infantis global population and to other *Salmonella* serovars is a grave public health concern (35).

## Conclusion

To conclude, the vast majority of *S*. Infantis isolates fall within eBG31 which is comprised of three clusters, a North American cluster (Cluster A), European cluster (Cluster C) and an ancestral but still extant global cluster (Cluster B). Isolates from Africa were genetically more diverse and distant from isolates from the other continents, further confirming previous work that identified a distinct population structure in South African *S*. Infantis. Using a One Health approach we were able to observe high levels of AMR in poultry and poultry products, highlighting the need to reduce the levels of this pathogen in poultry production premises and encouraging the development and use of a vaccine against *S*. Infantis in poultry. Finally, pESI was shown to be the major driver for AMR in the global *S*. Infantis population and presents a major threat to public health.

## Acknowledgements

We would like to thank Anaïs Painset in GBRU, UKHSA for their help with bioinformatics training. We are grateful to Heather Carleton from Centers for Disease Control and Prevention, USA, for sharing isolate metadata with us.

## Data availability

The Illumina FASTQ accessions for all the isolates are available in Table S1a. The ARIBA ResFinder, Plasmidfinder and gyrase results are in Table S1b, c and d. The eBG31 reference genome can be accessed under the accession CP070301.

