## Supplementary Information for "*Salmonella* Infantis, the emerging human multidrug resistant pathogen – a One Health perspective"

**Methods**

**Isolate selection**

Enterobase was searched on the 19^th^ of February 2018, and all isolates belonging to eBG31 with sequence data available in the short read archive (SRA) were included; due to the computational cost of including more sequences we chose to not include isolates after this cut-off (1). All UK Health Security Agency (UKHSA) eBG31 sequence data reported by 31.12.20 was downloaded from the UKHSA Pathogens BioProject at NCBI (PRJNA248792). Sequence data for 62 Animal and Plant Health Agency (APHA) *S*. Infantis isolates were downloaded from the SRA on 28.05.19.

DNA of *S*. Infantis isolates was shared for sequencing by the APHA, UKHSA and National Institute of Communicable Diseases (NICD), South Africa. The APHA isolates included had been collected for surveillance up until March 2018. Strains, isolated between 2000 and 2014, were selected from the UKHSA culture store; ensuring that there was an even distribution by time and, if applicable, continent of travel. Strains isolated between 2004 and 2016 were selected from the NICD culture store for sequencing.

The metadata available was used to stratify the isolates into the following source groups: human, food, poultry, poultry products, animal feed, other animals, environmental and unknown (Supplementary Table 1a). All clinical isolates and those with metadata on patient age or gender were classified as human. Isolates from chickens, ducks, turkeys and quail were categorised as poultry. All strains from poultry products, such as eggs and chicken meat, were grouped together. Isolates from all other animals were grouped into ‘other animals’ and non-poultry animal products and other food samples were grouped into ‘food’. All samples from animal feed were grouped and farm swabs, soil, water, air and sewage were designated environmental. Isolates missing source information to assign to these groups were classed as unknown. For the UKHSA isolates with travel history information, the continent of travel was used to designate the continent of isolate origin. The world map plot was generated with R (v.4.1.0) and the package ggplot2 (v.3.3.6) (2,3).

**Whole Genome Sequencing**

DNA from the NICD isolates was extracted using QIAamp DNA Mini Kit (Qiagen) and sequenced as described in Mattock *et al*., 2021 (4); libraries were prepared using either Illumina Nextera XT library preparation kits or using a custom library preparation method described in Rasheed *et al*., 2020 (5). Twelve NICD isolates not used in Mattock *et al.*, 2021 are included here and a library was prepared using one of the aforementioned methods and sequenced on an Illumina NextSeq 500.

The DNA of the UKHSA samples was extracted using the Qiasymphony DSP DNA Midi Kit (Qiagen), following a protocol described in Nair *et al*., 2016 (6). The historical UKHSA samples, described in Lee *et al*., 2021, were also sequenced using either Illumina Nextera XT or the custom library preparation methods (7). Two additional historical UKHSA samples were included and sequenced using the custom library preparation method. The contemporary UKHSA samples were prepared using the Illumina Nextera XT protocol and sequenced on an Illumina HiSeq 2500 instrument.

DNA from the APHA isolates was extracted using the MagMAX CORE Nucleic Acid Purification Kit (ThermoFisher) with a KingFisher Flex Purification System (ThermoFisher). The DNA was then sequenced using the custom library preparation method and an Illumina NextSeq 500.

**Phylogenetic analysis**

All the sequence data generated by UKHSA were trimmed using Trimmomatic (v0.27) with leading and trailing at <Q30 (8). All other sequence data were trimmed with Trimmomatic (v0.36) and the same options. Sequence type and eBG were determined for each isolate using Metric-Oriented Sequence Typer (MOST) (v.1.0) with a UKHSA *Salmonella* database (9,10). The quality of borderline sequence typing was assessed using Tablet (11).

Using the Cloud Infrastructure for Big Data Microbial Bioinformatics (12), each sample was mapped and variant called against the eBG31 reference CP070301 (7), using Snippy (v.4.6.0) with the options minfrac 0.9 and 30 (13). Due to the large number of sequences included, it was not computationally possible to produce a phylogeny including all of the isolates. Therefore, the SNP distance between all of the consensus FASTAs produced by Snippy was calculated using snp-dists (0.7) (<https://github.com/tseemann/snp-dists>) with the -m option to output in the molten format. Using MCL (v14-137) and the abc option, Markov clustering was performed on the pairwise SNP distance matrix with the following SNP distance thresholds: 0 SNPs, 5SNPs, 10SNPs, 25SNPs, 50SNPs, 100SNPs and 250SNPs (10,14). This resulted in clusters where each cluster member was less than ‘n’ SNPs from another member. The smallest SNP threshold that it was possible to generate a phylogeny with was 25SNPs; the first isolate in each 25SNP cluster was chosen to be the representative for that cluster.

The eBG31 reference genome, CP070301, was screened for prophages using PHASTER (7,15). Four complete prophages were identified and masked by Snippy when producing an alignment of the 25SNP cluster representatives. The whole genome alignment was used by Gubbins (v.2.4.1) to identify recombination which was then removed during core-SNP alignment generation (16). A core SNP maximum likelihood phylogeny of the 25SNP cluster representatives, excluding the reference isolate, was constructed using RaxML (v.8.2.12), rooted to its most ancestral node and annotated using iToL (17,18). Hierarchical Bayesian clustering within the phylogeny was identified using fastbaps (v.1.0.5) with ape (v.5.3) and R (v.3.4.1), with the optimised symmetric prior and k.init at 5 (19–21). Pairwise SNP distances were calculated using the snp-dists output, with self against self comparisons excluded.

The treedater relaxed clock test was performed to determine support for using a strict clock; the strict clock was deemed the best approach (22). Treedater (v.0.5.0) was used with a strict clock and the dates ranges of the isolates in each 25SNP cluster, excluding leaves with an unknown isolation date, to date the phylogeny. Phangorn (v.2.10.0) was used to identify the nodes where the clades diverged (23).

**AMR and plasmid determination**

ARIBA (v.2.10.1) was used with the ResFinder and Plasmidfinder databases (downloaded 11.04.19) to identify AMR determinants and plasmids (24–26). A database of *gyrA*, *gyrB*, *parC* and *parE*, from the *S*. Typhimurium LT2 reference, was used with ARIBA to identify mutations within the Quinolone Resistance Determining Regions (QRDR). MDR, resistance to three or more antimicrobial classes, was calculated using AMR genes identified by ARIBA and mutations in QRDRs. As aminoglycoside resistance in *Salmonella* is rarely conferred by *aac(6’)-Iaa* it was excluded from all calculations (27). pESI presence was determined by mapping each sequence against a pseudomolecule of the eBG31 reference (CP070301) and pESI contigs (ASRF01000099-ASRF01000108) with SMALT (v.0.7.6) and calculating coverage with Samtools (v.1.5) (28,29). Heatmaps were generated using phytools (v.0.6) and data.table (v.1.11.8) (30,31).

The proportion of isolates containing each AMR determinant and plasmid from each metadata group was calculated in R (v.4.1.0) with the packages janitor (v.2.1.0), magrittr (v.2.0.1) and tidyr (v.1.1.3) (32–34). The heatmap was created in R with ape and gplots (v3.1.3) (35).

**Supplementary Tables**

**Table S1 *S*. Infantis metadata, ARIBA ResFinder, Plasmidfinder and gyrase results**

a) Metadata of the 5284 *S*. Infantis isolates with their isolation date, country, continent, source group, all other available source metadata, fastbaps cluster and ST
b) AMR gene presence/absence as determined by ARIBA with the ResFinder database
c) Plasmid presence/absence as determined by ARIBA with the Plasmidfinder database and pESI presence/absence
d) Mutations in the QRDRs

**Supplementary Figures**

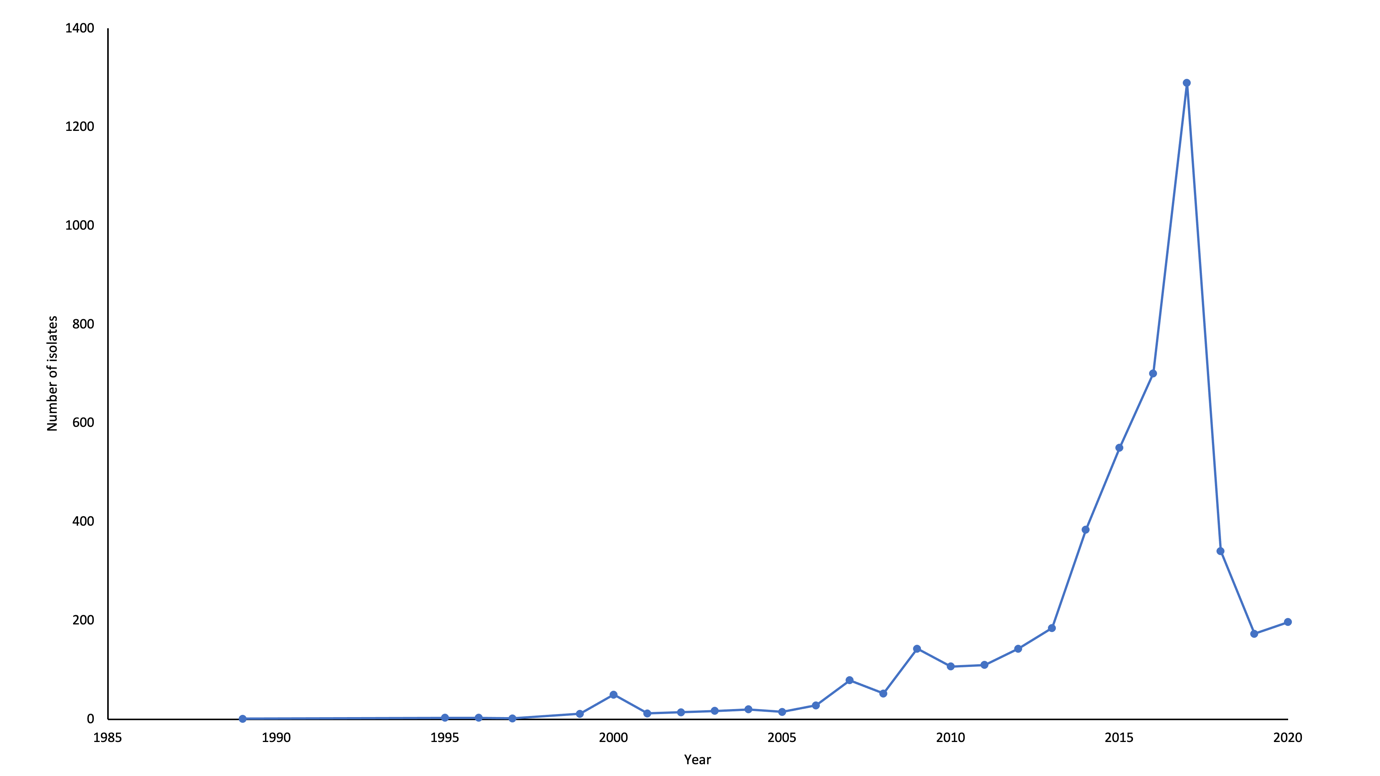
**Figure S1 Frequency of *S*. Infantis each year**

Number of *S*. Infantis strains included in the dataset from 1989 to 2020

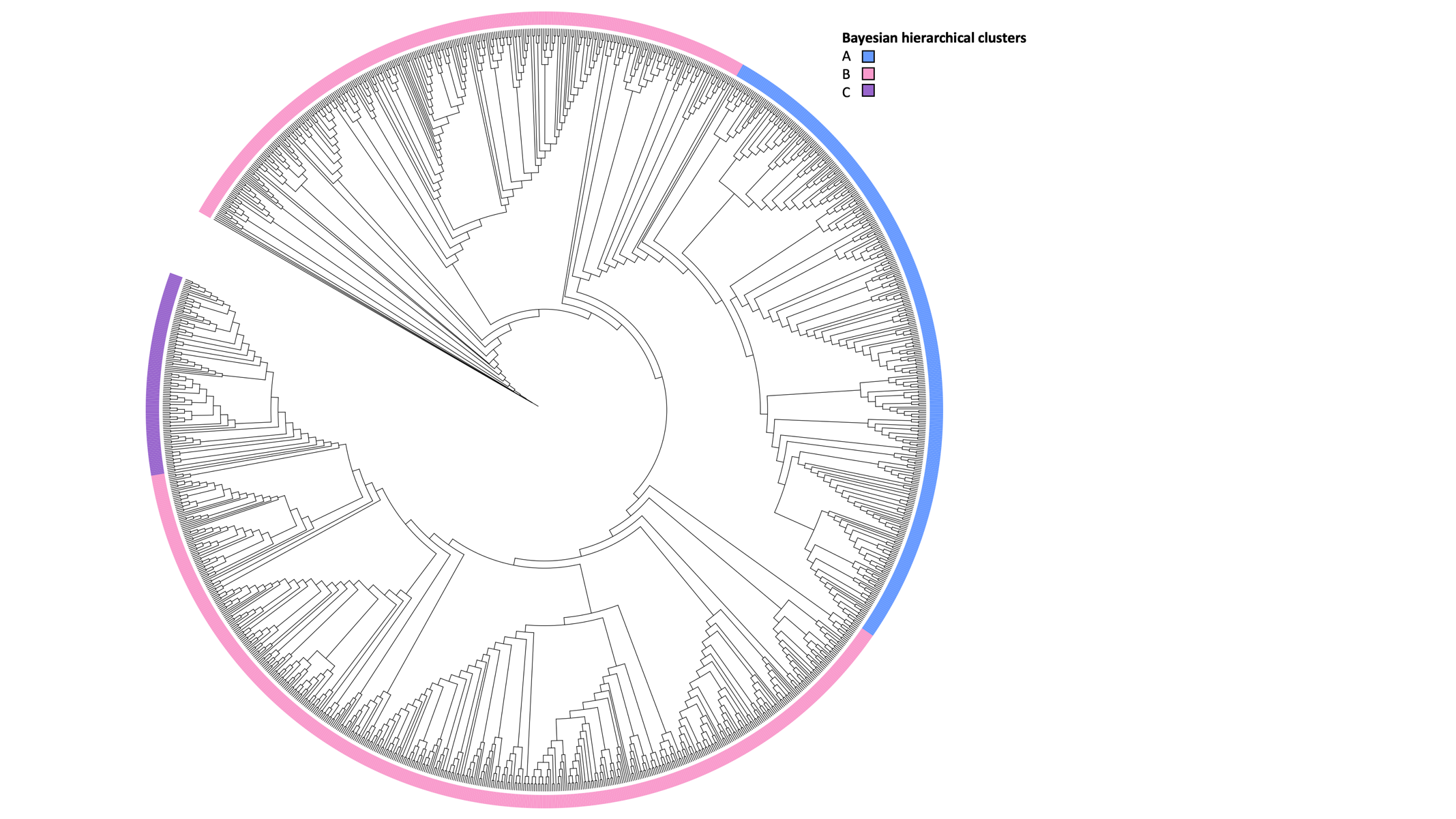

**Figure S2 Maximum likelihood cladogram of *S*. Infantis**

Core SNP maximum likelihood cladogram of 1288 representatives of 5283 *S*. Infantis isolates. The ring around the cladogram is annotated with the Bayesian hierarchical clusters found by fastbaps.

Bayesian hierarchical clusters: A, 348 representatives of 1624 isolates; B, 831 representatives of 3283 isolates and C 109 representatives of 376 isolates

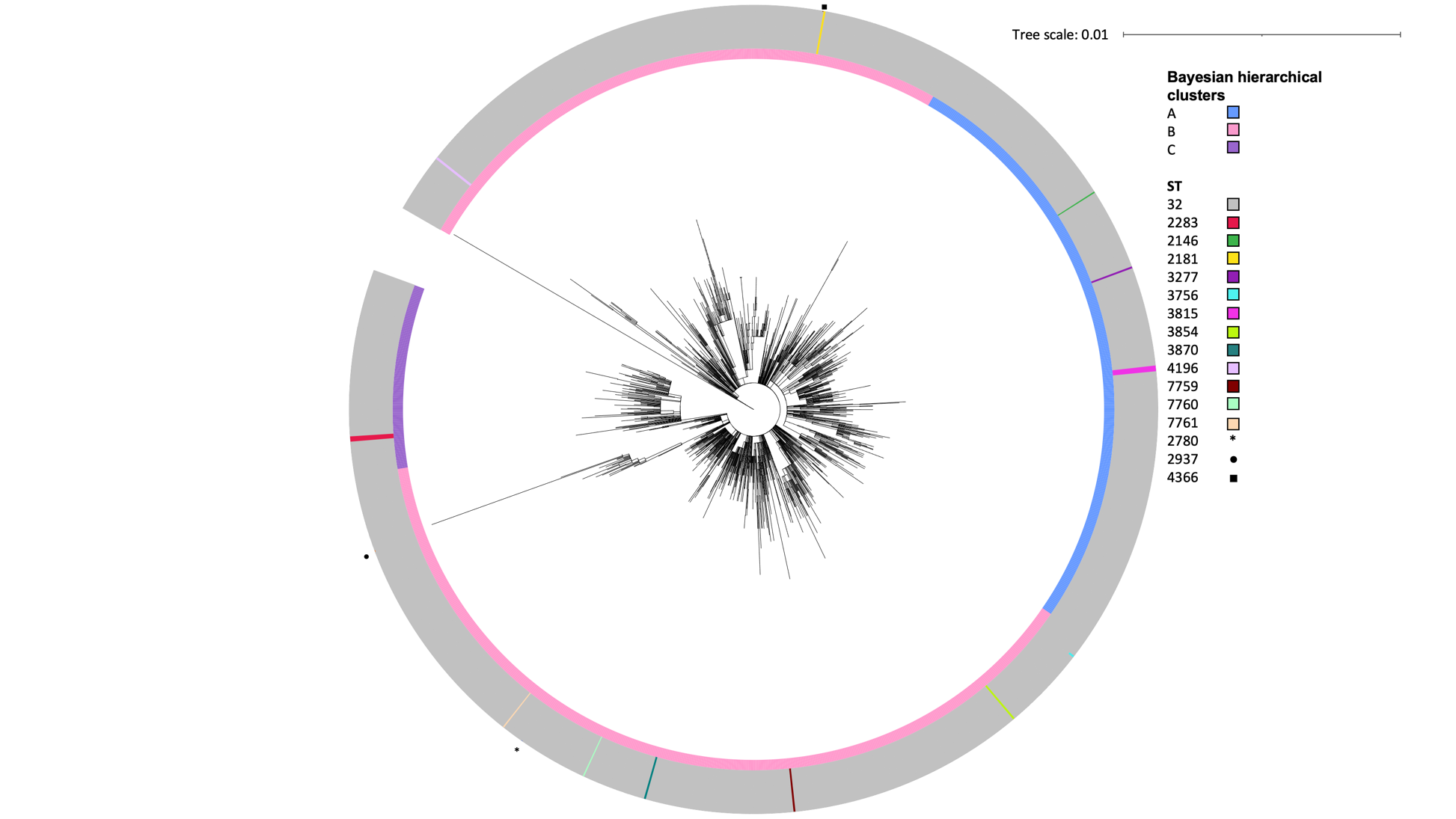

**Figure S3 Maximum likelihood phylogeny of *S*. Infantis with ST**

Core SNP maximum likelihood phylogeny of 1288 representatives of 5283 *S*. Infantis isolates. The clusters in the phylogeny are annotated with the Bayesian hierarchical clusters found by fastbaps. The outer ring shows the percentage of isolates in each 25SNP cluster that belonged to each ST.

Bayesian hierarchical clusters: A, 348 representatives of 1624 isolates; B, 831 representatives of 3283 isolates and C 109 representatives of 376 isolates

STs: 32 (n=5204), 2283 (n=36), 2780 (n=1), 2181 (n=1), 3756 (n=2), 3277 (n=1), 2146 (n=26), 2937 (n=2), 3815 (n=3), 3870 (n=1), 3854 (n=1), 4196 (n=1), 4366 (n=1), 7759 (n=1), 7760 (n=1), 7761 (n=1)

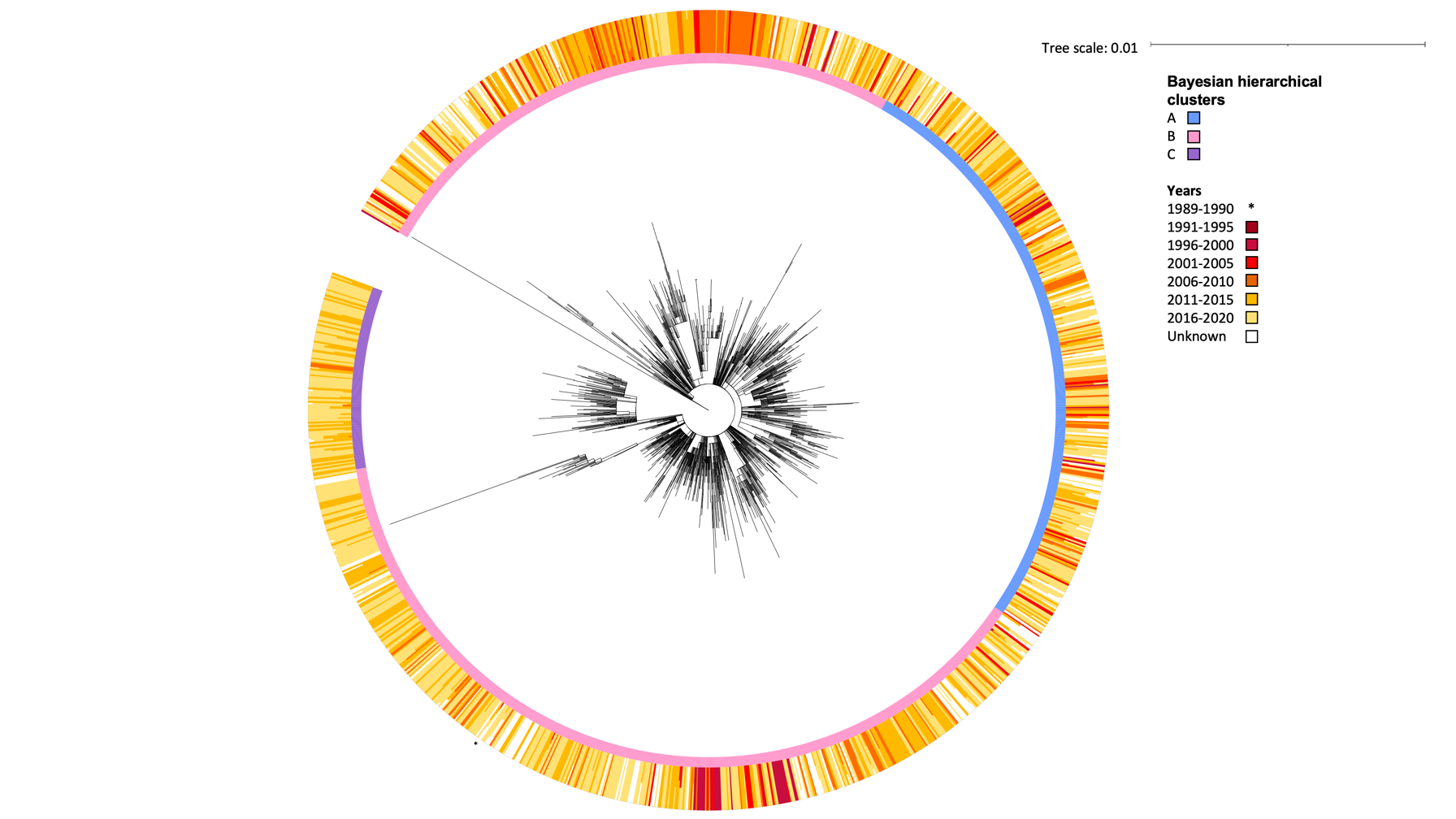

**Figure S4 Maximum likelihood phylogeny of *S*. Infantis with isolation year**

Core SNP maximum likelihood phylogeny of 1288 representatives of 5283 *S*. Infantis isolates. The clusters in the phylogeny are annotated with the Bayesian hierarchical clusters found by fastbaps. The outer ring shows the percentage of isolates in each 25SNP cluster that were from each year group.

Bayesian hierarchical clusters: A, 348 representatives of 1624 isolates; B, 831 representatives of 3283 isolates and C 109 representatives of 376 isolates

Years: 1989-1990 (n=1), 1991-1995 (n=3), 1996-2000 (n=66), 2001-2005 (n=78), 2006-2010 (n=409), 2011-2015 (n=1371), 2016-2020 (n=2702), unknown (n=653)

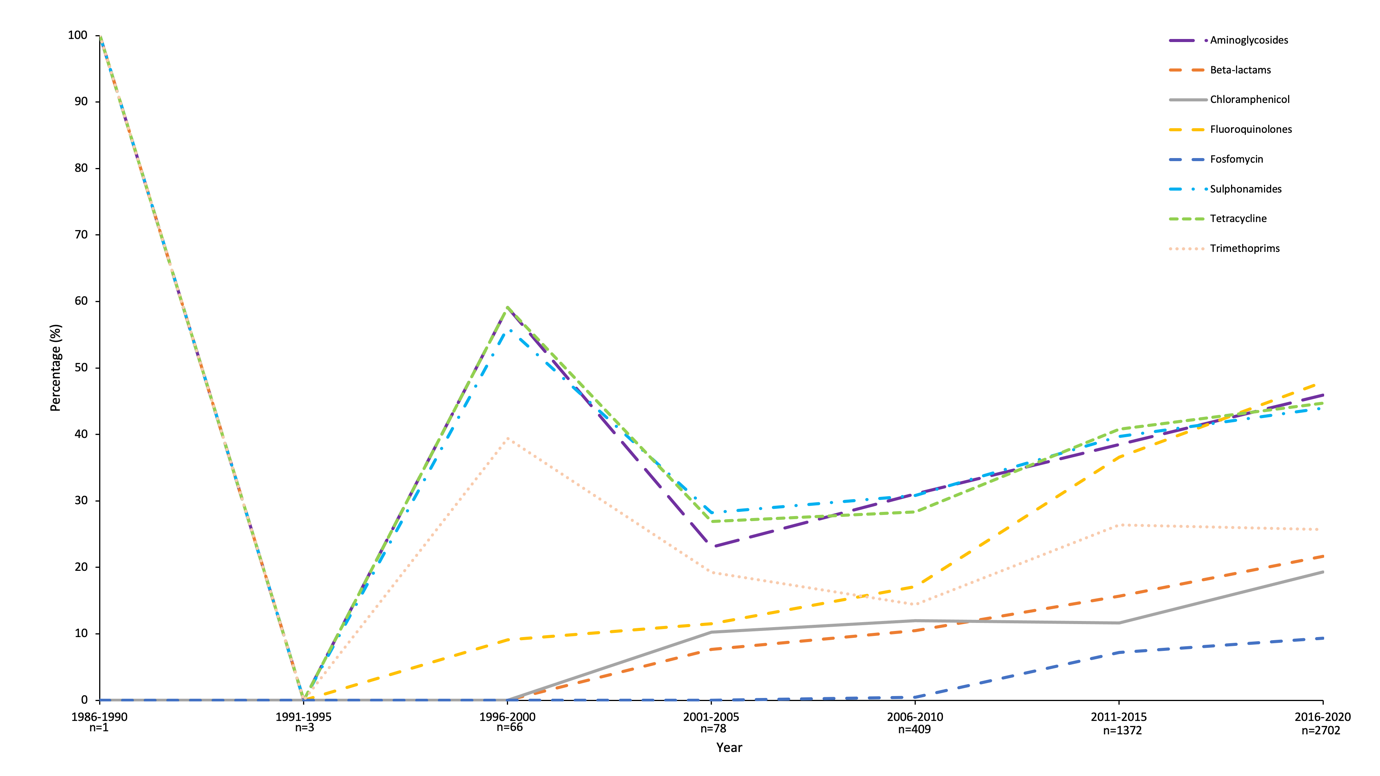

**Figure S5 AMR in *S*. Infantis each year**

Proportion of *S*. Infantis isolates from each year with genes conferring resistance to the antimicrobial classes that had resistance in >5% of isolates: aminoglycosides, beta-lactams, chloramphenicol, fluoroquinolones, sulphonamides, tetracyclines and trimethoprim.

Years: 1989-1990 (n=1), 1991-1995 (n=3), 1996-2000 (n=66), 2001-2005 (n=78), 2006-2010 (n=409), 2011-2015 (n=1372), 2016-2020 (n=2702), unknown (n=653)
